# Assigning functionality to cysteines by base editing of cancer dependency genes

**DOI:** 10.1101/2022.11.17.516964

**Authors:** Haoxin Li, Jarrett R. Remsberg, Sang Joon Won, Kevin T. Zhao, Tony P. Huang, Bingwen Lu, Gabriel M. Simon, David R. Liu, Benjamin F. Cravatt

## Abstract

Chemical probes are lacking for most human proteins. Covalent chemistry represents an attractive strategy for expanding the ligandability of the proteome, and chemical proteomics has revealed numerous electrophile-reactive cysteines on diverse proteins. Determining which of these covalent binding events impact protein function, however, remains challenging. Here, we describe a base-editing strategy to infer the functionality of cysteines by quantifying the impact of their missense mutation on cell proliferation. We show that the resulting atlas, which covers >13,800 cysteines on >1,750 cancer dependency proteins, correctly predicts the essentiality of cysteines targeted by cancer therapeutics and, when integrated with chemical proteomic data, identifies essential, ligandable cysteines on >110 cancer dependency proteins. We further demonstrate how measurements of reactivity in native versus denatured proteomes can discriminate essential cysteines amenable to chemical modification from those buried in protein structures, providing a valuable resource to prioritize the pursuit of small-molecule probes with high function-perturbing potential.

## Introduction

Small molecules are powerful tools for studying the functions of proteins in biological systems and can serve as starting points for therapeutics^1^. Advanced chemical probes and drugs frequently target established small-molecule binding pockets in proteins, such as the active sites of enzymes (e.g., ATP-competitive inhibitors of kinases^2^) or ligand-binding pockets of receptors (e.g., antagonists of androgen and estrogen receptors^3^). Many protein types, however, are not known to bind endogenous small molecules and are consequently more difficult to gauge in terms of their potential for targeting by chemical probes. The timeliness and importance of this problem are accentuated by the output of modern large-scale sequencing and genetic screening efforts, which are identifying a broad array of human disease-relevant genes that code for protein types such as DNA/RNA-binding or adaptor/scaffolding proteins^4–6^ that have historically been challenging to target with small molecules.

Several platforms for the discovery of small-molecule binders of proteins have recently been introduced as a means to expand the ligandability of the human proteome^7^. These ‘binding-first’ technologies, which include fragment-based screening^8^, DNA-encoded libraries^9^, and chemical proteomic methods such as activity-based protein profiling (ABPP)^10,11^, have led to the discovery of small-molecule ligands for diverse arrays of proteins. Nonetheless, whether such ligands affect the functions of protein targets often remains unclear and can be challenging to determine experimentally, especially for proteins with poorly characterized biochemical or cellular activities. In cases where newly discovered small molecule-protein interactions have been shown to be functional, it is notable that they frequently act by allosteric mechanisms^12,13^, underscoring the diverse and unanticipated ways that chemical probes can modulate the activity of proteins.

Among binding-first approaches, the chemical proteomic analysis of electrophilic small molecules has demonstrated considerable potential for ligand discovery^10,11,14–20^. Advantages of this approach include the deployment of covalent chemistry^19,21–23^, which can address shallow and dynamic pockets on protein surfaces that are less amenable to binding reversibly to small molecules, as well as provide a selectivity filter in the form of targeting isotype-restricted nucleophilic residues (e.g., cysteines) within paralogous proteins, as have been demonstrated by several recent cancer therapeutics (e.g., osimertinib for EGFR_C797^24,25^, sotorasib for KRAS_G12C^26,27^). Additionally, by evaluating small molecules directly in native biological systems, chemical proteomics methods such as ABPP can identify cryptic ligandable pockets regulating aspects of protein function that are difficult to discern with purified proteins or protein domains^17,28,29^.

The human genome encodes over 200,000 cysteines distributed across virtually all proteins. A growing, but still modest proportion of these cysteines has been shown to interact with electrophilic small molecules in ABPP experiments^10,16–19,21^, and, even in these cases, the impact of electrophile-cysteine interactions on protein function remains mostly unknown. This latter question is important for prioritizing the pursuit of advanced chemical probes and, ultimately, drugs that act through covalent modification of cysteines.

Here, we introduce an integrated base editing and chemical proteomic strategy for globally assessing the functionality and ligandability of cysteine residues in the context of cancer dependency proteins as defined by the Cancer Dependency Map^5^. By specifically altering the codons encoding cysteines and neighboring residues using programmable base editors comprising Cas9 nickases fused to cytidine deaminases or laboratory-evolved deoxyadenosine deaminases^30,31^ and recording the impact of these missense mutations on cancer cell proliferation, we discover evidence of essentiality for 1,359 cysteines (among >13,800 cysteines evaluated) on diverse classes of cancer dependency proteins. Combining these data with chemical proteomic profiles of fragment electrophile-cysteine interactions in cancer cells furnished a prioritized list of >110 essential, ligandable cysteines. These cysteines include validated sites of covalent drug action and many additional residues on proteins with Common Essential or Strongly Selective designations in the Cancer Dependency Map^5^. Finally, we present a general strategy to categorize essential, but as-of-yet unliganded cysteines based on their chemical reactivity in native versus denatured proteomes, which can be used to distinguish buried residues likely performing structural roles in proteins from those that are solvent accessible with future ligandability potential. The platforms described herein thus offer residue-level functional annotation and small-molecule reactivity maps to guide the ongoing and future pursuit of covalent chemical probes and drugs.

## Results

### Base editing assigns essentiality to cysteines targeted by anti-cancer drugs

Base editing enzymes catalyze nucleotide substitutions at targeted genomic DNA sites using programmable DNA binding proteins fused to cytidine or deoxyadenosine deaminases^32,33^. Within an “editing window”, deoxyadenosine deaminases in adenine base editors (ABEs) convert an A•T base pair into a G•C base pair^30^ and cytidine deaminases in cytosine base editors (CBEs) convert a C•G base pair into a T•A base pair^31^. For base editing to inform the essentiality of cysteines in proteins that are required for cancer cell proliferation, the mutagenic paths afforded by ABEs (cysteine-to-arginine; **Fig. 1a**) and CBEs (cysteine-to-tyrosine; **Fig. 1a**) would need to have a good probability of altering protein function. We explored this concept by categorizing 65,000 missense mutations with confirmed or potential pathogenic phenotypes based on the ClinVar database^34^ and found 3,531 cases where cysteine was mutated, of which 18.8% and 23.3% represented conversions to arginine and tyrosine, respectively (**Fig. 1b**). These frequencies were much larger than the pathogenic missense mutations associated with converting cysteine to serine, phenylalanine, or glycine (~5-10%), despite equivalent or greater codon availability for these mutational paths (**Fig. 1b**). Importantly, commonly used ABEs and CBEs are only capable of converting cysteine to arginine and tyrosine, respectively, as other base-editing products catalyzed by these enzymes yield synonymous mutations (**Fig. 1a** and **Extended Data Fig. 1a, b**). Taken together, these findings suggested that base editing should serve as a suitable strategy to evaluate the functional consequences of mutating cysteines.

**Fig. 1.**
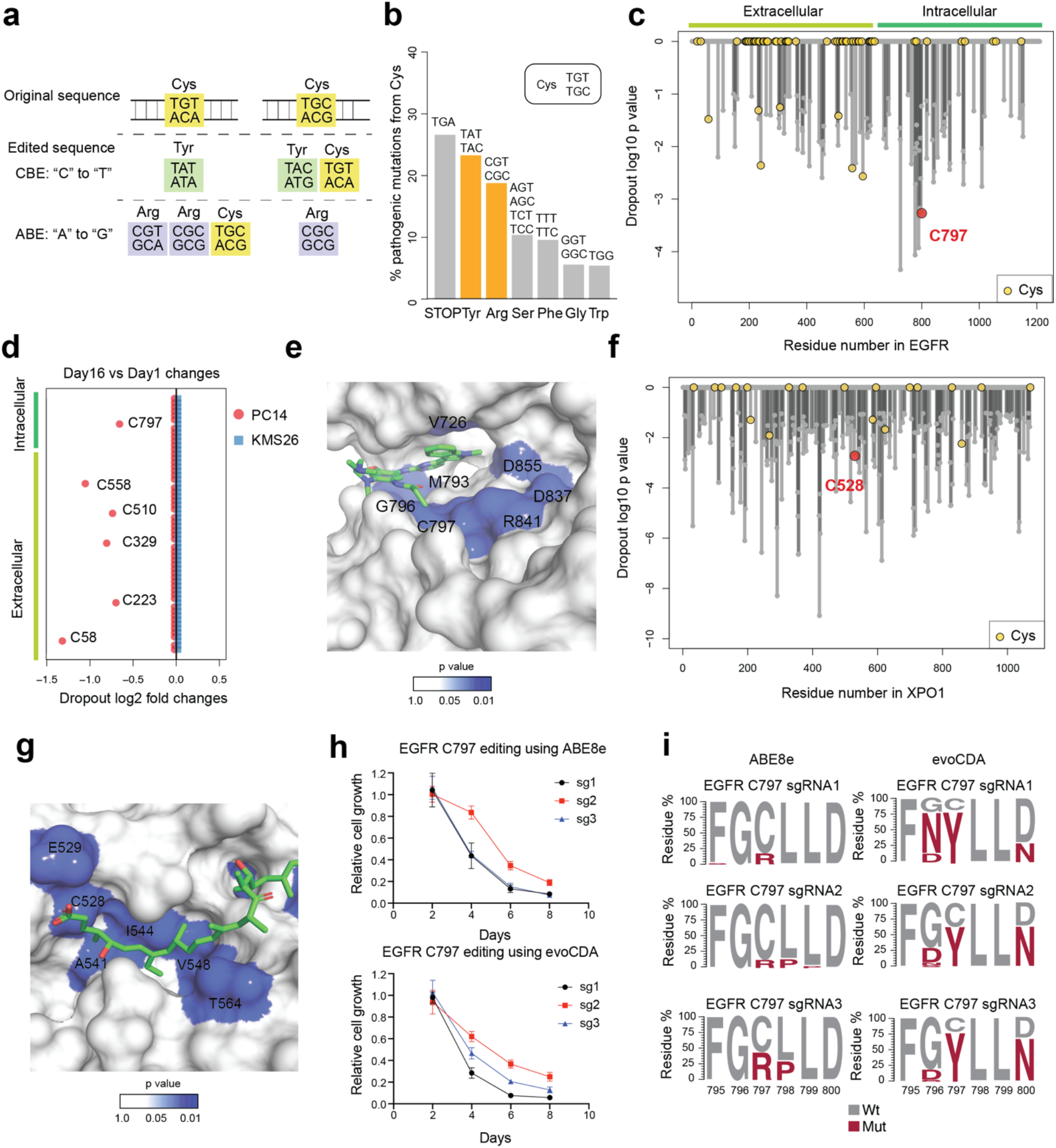
Base editing assigns essentiality to cysteines targeted by anti-cancer drugs. **a**, Codon analysis of cysteine mutagenesis patterns generated by Adenine Base Editors (ABE) or Cytosine Base Editors (CBE). **b**, Barplot showing the distribution of pathogenic missense mutations of cysteine residues (N = 3531) from the ClinVar database. **c**, Waterfall plot showing the dropout significance calculated per EGFR residue between day 16 vs day 1 from saturated scanning experiments. C797 is marked as a red circle, and other cysteine residues are highlighted as yellow circles. Data represent average values of two independent experiments. P values were calculated using Student’s t test**. d**, Dot plot comparing the day 16 vs day 1 log2 fold dropout changes (LFC) for sgRNAs targeting cysteine residues in EGFR in PC14 and in KMS26 cells. Cutoff: LFC < −0.4; p < 0.05. Data represent average values of two independent experiments. **e**, Structure of the EGFR kinase domain bound to osimertinib (PDB: 6JXT) with residues colored from white to blue to indicate increased dropout significance. **f**, Waterfall plot showing the dropout significance calculated per XPO1 residue between day 16 vs day 1 from saturated scanning experiments. C528 is marked as a red circle, and other cysteine residues are highlighted as yellow circles. Data represent average values of two independent experiments. P values were calculated using Student’s t test**. g**, Structure of the XPO1 cargo-binding cleft in association with the natural product Leptomycin B (PDB: 6TVO) with residues colored from white to blue to indicate increased dropout significance. **h**, Proliferation graphs for PC14 cells with EGFR_C797 editing in an arrayed format measured by CellTiterGlo. The relative growth was normalized to 100% on day 2. Data represent average values ± SEM for four independent experiments. **i**, EGFR amino acid mutagenesis generated by different sgRNAs and the indicated base editors in PC14 cells. The relative residue frequency five days after lentivirus infection was quantified using targeted genomic PCR and amplicon sequencing. Data represent average values of two independent experiments.

We selected ABE8e and evoCDA in conjunction with SpCas9-NG to recognize an “NG” protospacer adjacent motif (PAM) specificity^35–37^ as the base editors for our study because they provided a combination of high editing efficiency, broad genome-wide targeting scope, and suitable editing window sizes (~5 nucleotides (nt) for ABE8e and ~10 nt for evoCDA) in comparison to other base editors. The 5-10 nt editing windows allowed for missense mutation of a greater number of cysteines within reach of a PAM site, but also introduced the potential to edit neighboring residues. We considered this approach to offer a reasonable balance of specificity and generality and anticipated that edits at nearby residues could provide useful information on the essentiality of the local environment surrounding a ligandable cysteine.

We first evaluated the performance of our cysteine editing protocol with two established cancer dependency proteins – EGFR and XPO1 – that are targets of cysteine-directed covalent drugs. Specifically, C797 of EGFR is engaged by covalent inhibitors such as osimertinib to treat non-small cancer lung cancer (NSCLC)^34^, and C528 of XPO1 is engaged by selinexor, a first-in-class inhibitor of this nuclear export protein to treat multiple myeloma^38^. We designed a pooled base-editing library containing >3,000 sgRNAs representing a mix of ABE8e and evoCDA sites that covered most residues (“saturated scan”) in EGFR and XPO1 and delivered this library to the EGFR-dependent NSCLC cancer cell line PC14 and the multiple myeloma cell line KMS26 using lentiviral vectors (**Extended Data Fig. 1c**). We then allowed the cells to grow for 16 days and performed targeted amplicon sequencing of the sgRNA cassette region on days 1 and 16 to obtain relative sgRNA frequency changes associated with cancer cell proliferation. The dropout values for all sgRNAs predicted to edit the same residue were averaged to give a significance value of residue-level essentiality (**Supplementary Table 1**). Encouragingly, this analysis identified EGFR_C797 among the most essential residues and the top dropout among all intracellular cysteines in EGFR in PC14 cells (**Fig. 1c, d**). Other essential cysteines in EGFR were extracellular residues involved in structural disulfide bonds (**Fig. 1c, d**). We did not observe essentiality for EGFR_C797 or, in general, other EGFR residues in KMS26 cells (**Fig. 1d**), consistent with this cell line being independent of EGFR for growth. The preferential essentiality of C797 relative to other intracellular cysteines in EGFR was also observed when individual data sets from ABE8e or evoCDA were analyzed (**Extended Data Fig. 1d**). Interestingly, we also found evidence of essentiality for other residues located near C797 in and around the kinase catalytic pocket, including V726, M793, G796, D837, R841, and D855 (**Fig. 1e**). Similar findings emerged for XPO1, a common essential protein involved in nuclear export^39^, where the aggregated data using both model cell lines suggested that C528 and additional residues (e.g., E529, A541, I544, V548, T564) in the XPO1 substrate/selinexor-binding cleft displayed significant dropout values (**Fig. 1f, g**).

We validated the pooled screening results by individual cloning and gene editing of six selected EGFR_C797-targeting sgRNAs in an arrayed format. We observed that all six sgRNAs caused prominent decreases in PC14 proliferation (**Fig. 1h**), and targeted genomic sequencing revealed editing of EGFR_C797 and the region flanking this residue (**Fig. 1i**). Consistent with the predicted editing specificities and windows, ABE generated a C797R mutation with or without the presence of an L798P mutation, while CBE produced a C797Y mutation in the presence of G796N/D and D800N mutations (**Fig. 1i**).

These proof-of-principle experiments with established cancer dependency proteins supported that our strategy combining base editing with pooled cell proliferation assays can create sufficient single or composite mutations at and around cysteines targeted by covalent small molecules to reveal the functionality of these residues and the druggable pockets where they are located.

### Global analysis of essential cysteines in cancer dependency proteins

We next sought to globally assess the functionality of cysteines in a broad set of cancer dependency proteins by base editing combined with pooled cell proliferation assays. We designed ~50,000 sgRNAs using both ABE and CBE editors that covered >13,800 cysteines from >1,750 cancer dependency proteins. Of these proteins, ~270 were defined as Strongly Selective in the Cancer Dependency Map, reflecting a restricted dependency relationship with a subset of the cancer cell line panel, and another ~1500 were defined as Common Essential to indicate their general requirement for the growth of most cancer cell lines (**Fig. 2a** and **Extended Data Fig. 2a**). The sgRNA library contained at least one sgRNA that satisfied the NG PAM requirements of the ABE or CBE editors for about 70% of all cysteines in the cancer dependency proteins (**Extended Data Fig. 2b**). Theses sgRNAs were screened as a pooled library, along with ~1,200 non-targeted control sgRNAs, in separate experiments performed in the PC14 and KMS26 cell lines, and, after 16 days, sgRNA frequency changes compared to day 1 were quantified by targeted sequencing (**Fig. 2b**).

**Fig. 2.**
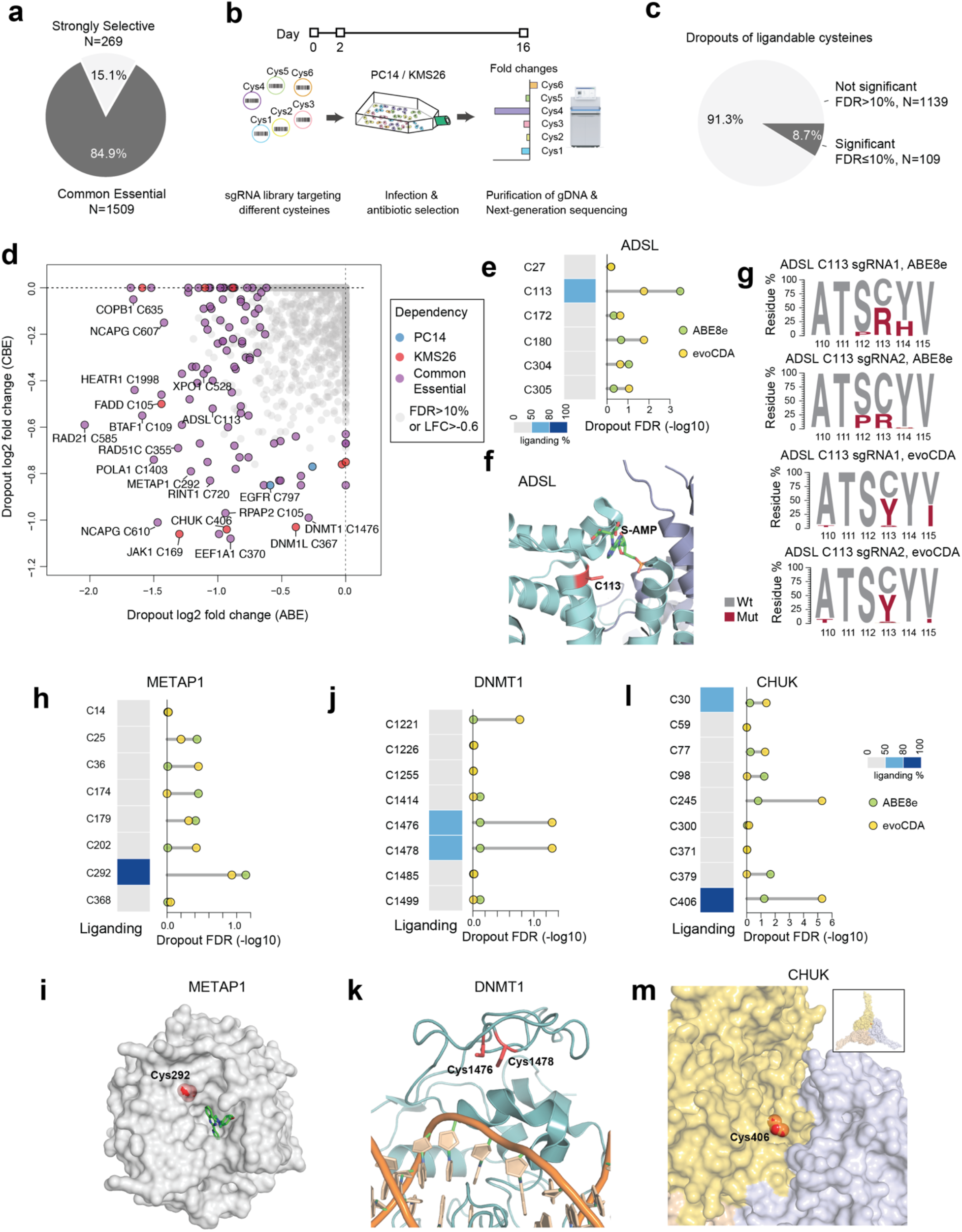
Global discovery of essential cysteines in cancer dependency proteins. **a**, Pie chart showing the proportion of Strongly Selective and Common Essential proteins (as assigned by the Cancer Dependency Map) evaluated in the cysteine-directed, base editing dropout screen. **b**, Schematic showing the workflow for determining cysteine essentiality by base editing dropout screens performed in human cancer cell lines. In brief, spin-infection with pooled lentivirus libraries delivering base editors and sgRNAs was performed on day 0 and puromycin was added on day 2. Cells were cultured for two more weeks and the pellets were collected for next-generation sequencing. Independent experiments were performed in duplicates and dropout signals were averaged for each base-edited cysteine. **c**, Pie chart showing the proportion of ligandable cysteines (as assigned by chemical proteomic experiments with electrophilic fragments^10,22,32^ and data generated herein (**Supplementary Table 2**)) with or without significant dropouts in gene editing screens (FDR cutoff: 10%, LFC cutoff: −0.6). **d**, Overview of essential, ligandable cysteines identified in gene editing dropout screens. The log2 fold change (LFC) of each cysteine is calculated by comparing the frequency of corresponding sgRNAs between day 16 vs day 1 (FDR cutoff: 10%, LFC cutoff: −0.6). The p values were estimated by comparing observed dropout LFC to the dropout LFC of non-targeting sgRNAs. For cysteines from Common Essential proteins, the average dropout values of both model cell lines were used in the calculations, and, for cysteines from Strongly Selective proteins, either PC14 (blue points) or KMS26 (red points) dropout data were used. **e**, The fragment electrophile ligandability profile (left) and significance of dropout (right) for cysteines in ADSL. **f**, Structure of ADSL in complex with its substrate adenylosuccinate (PDB: 2VD6). The side chain for essential C113 is highlighted in red. **g**, ADSL amino acid mutagenesis generated by different sgRNAs and the indicated base editors in PC14 cells. The relative residue frequency five days after lentivirus infection was quantified using targeted genomic PCR and amplicon sequencing. Data represent average values of two independent experiments. **h**, The fragment electrophile ligandability profile (left) and significance of dropout (right) for cysteines in METAP1. **i**, Structure of METAP1 in complex with the non-covalent pyridinylquinazoline inhibitor FZ1 (PDB: 4IU6). The side chain of essential C292 is highlighted in red. **j**, The fragment electrophile ligandability profile (left) and significance of dropout (right) for cysteines in DNMT1. Note that C1476 and C1478 are on the same tryptic peptide, so their ligandability cannot be distinguished by cysteine-directed MS-ABPP. **k**, Structure of DNMT1 in complex with DNA (PDB: 7SFG). The side chains of essential cysteines C1476 and C1478 are highlighted in red. **l**, The fragment electrophile ligandability profile (left) and significance of dropout (right) for cysteines in CHUK. **m**, Structure of CHUK showing the homo-oligomerization surface (PDB: 5EBZ). Each oligomer is shown in yellow, pink, or light blue. The side chain of essential C406 is highlighted in red.

Using parameters of gene-level effect^5^ (gene-level dependency score < - 0.4) and residue-level dropout (log2 fold changes (LFC) < −0.6; false discovery rate, FDR < 10%), we identified a total of 1,359 essential cysteines across 725 proteins (**Supplementary Table 2**). This % hit rate for essential cysteines (9.8% of 13,872 total screened cysteines) was much higher than the % hit rate observed with non-targeted sgRNAs (1.9%, **Extended Data Fig. 2c**). We then cross-referenced these results with chemical proteomic maps of cysteine engagement by broadly reactive fragment electrophiles (derived from past work^10,16,17^ supplemented by additional cysteine-directed ABPP experiments performed herein (**Supplementary Table 2**)), which yielded 109 cysteines showing combined features of essentiality and ligandability (**Fig. 2c** and **Supplementary Table 2**). A much larger set of ligandable cysteines (1,139 in total) found in the cancer dependency proteins were assigned as nonessential from the base-editing screens (**Fig. 2c** and **Supplementary Table 2**), suggesting only a limited relationship between cysteine ligandability and functionality, at least as determined by base editing-mediated mutagenesis analyzed in cell proliferation assays (see Discussion section). Essential, ligandable cysteines were distributed into sub-groups showing selective effects in PC14 or KMS26 cells, or in both cell lines (**Fig. 2d**), and this distribution generally matched that predicted from gene-level disruption (**Extended Data Fig. 2d-g**)^5^. The pooled screening strategy, for instance, rediscovered the selective essentiality of EGFR_C797 in PC14 cells and the common essentiality of XPO1_C528 in both PC14 and KMS26 cells (**Fig. 2d**).

Among the newly discovered essential cysteines were several intriguing cases. For example, the Common Essential metabolic enzyme adenylosuccinate lyase (ADSL), which is involved in *de novo* AMP synthesis, harbored a single ligandable cysteine C113 that exhibited significant dropout using either the ABE or CBE editor, and these effects were generally greater than those observed with other edited cysteines in ADSL (**Fig. 2e**). Structural analysis of ADSL showed that C113 is located in the substrate (adenylosuccinate) binding pocket (**Fig. 2f**), providing a rationale for the deleterious functional impact of mutating this residue. Arrayed targeted genomic sequencing of editing outcomes associated with each sgRNA in transduced cells revealed that the ABE created the expected C113R mutation along with S112P or Y114H mutations, while the CBE produced the expected C113Y mutation with or without the additional presence of a V115I mutation (**Fig. 2g**). A ligandable, active-site cysteine (C292) in methionine aminopeptidase 1 (METAP1) was also found to be essential for cancer cell growth (**Fig. 2h**), and this residue is in close proximity to the binding pocket for a reversible METAP1 inhibitor that suppresses initiator methionine processing on newly translated proteins^40^ (**Fig. 2i**).

Other essential, ligandable cysteines were found at protein-DNA and protein-protein interfaces. For instance, the essential, ligandable cysteines in DNMT1 (C1476 and C1478; **Fig. 2j**) – a Common Essential methyltransferase that maintains genome methylation patterns on newly synthesized hemimethylated DNA – are located distal to the active site of this enzyme in proximity to interactions with DNA (**Fig. 2k**). Similarly, an essential, ligandable cysteine (C406) in CHUK (IKK1/a) (**Fig. 2l**) – a serine kinase with a Strongly Selective dependency designation that regulates both canonical and non-canonical NF-κB signaling^41^ – resides in a pocket near a homo-oligomerization interface (**Fig. 2m**) reported to be critical for IKK1/a-dependent cellular processing of p100 to p52, a hallmark of non-canonical NF-kB signaling^42^.

These initial findings show how the integration of gene editing and chemical proteomic data can identify essential, ligandable cysteines required for cancer cell growth.

### Saturated local editing of ligandable cysteines in Strongly Selective cancer dependencies

While we found that the PC14 and KMS26 cell lines showed dependency on a substantial portion of Common Essential proteins (1,583 of 2,664 total such proteins defined in the Cancer Dependency Map), these cell lines required far fewer Strongly Selective proteins for growth (269 of 3,082 total in the Cancer Dependency Map)^5^. This is an expected outcome, as essential proteins receive the designation of Strongly Selective based on showing restricted impact on the growth of subsets of cancer lines, often reflecting lineage or mutational relationships^5^. Considering that many of the most compelling therapeutic targets in cancer (e.g., AR, BRAF, EGFR, ESR1, KRAS) fall into the Strongly Selective category, we devised an alternative strategy to expand our analysis of essential cysteines for this group of proteins. We specifically adopted a pooled dropout screening approach to enable parallel analysis of essential cysteines across twelve human cancer lines from nine different tissues of origin. This cell line panel together captured ~700 Strongly Selective proteins, of which ~30 possessed one or more ligandable cysteines (**Fig. 3a** and **Supplementary Table 3**). We focused our pooled dropout screen on these ligandable cysteines and expanded the targeting editing window to include three residues on either side of each cysteine, which we anticipated would increase our probability of obtaining high-efficiency editing events for assessing the essentially of the local cysteine region. This adaptation was made in response to finding that the editing efficiency for a substantial proportion of cysteines targeted in our initial screen (20% and 46% for CBE- and ABE-editing events, respectively) was < 20% (**Extended Data Fig. 3a, b**), which pointed to the possibility of overlooking the functional impact of ligandable cysteines due to inefficient editing, especially for sites targetable by very few sgRNAs. Finally, we noted that an additional advantage of the multiplexed pooled dropout screen is that individual cell lines should serve as cross-validating internal controls, where essential cysteines in Strongly Selective proteins would be expected to only impair the growth of the subset of cell lines in the panel that are dependent on that protein (**Fig. 3b**).

**Fig. 3.**
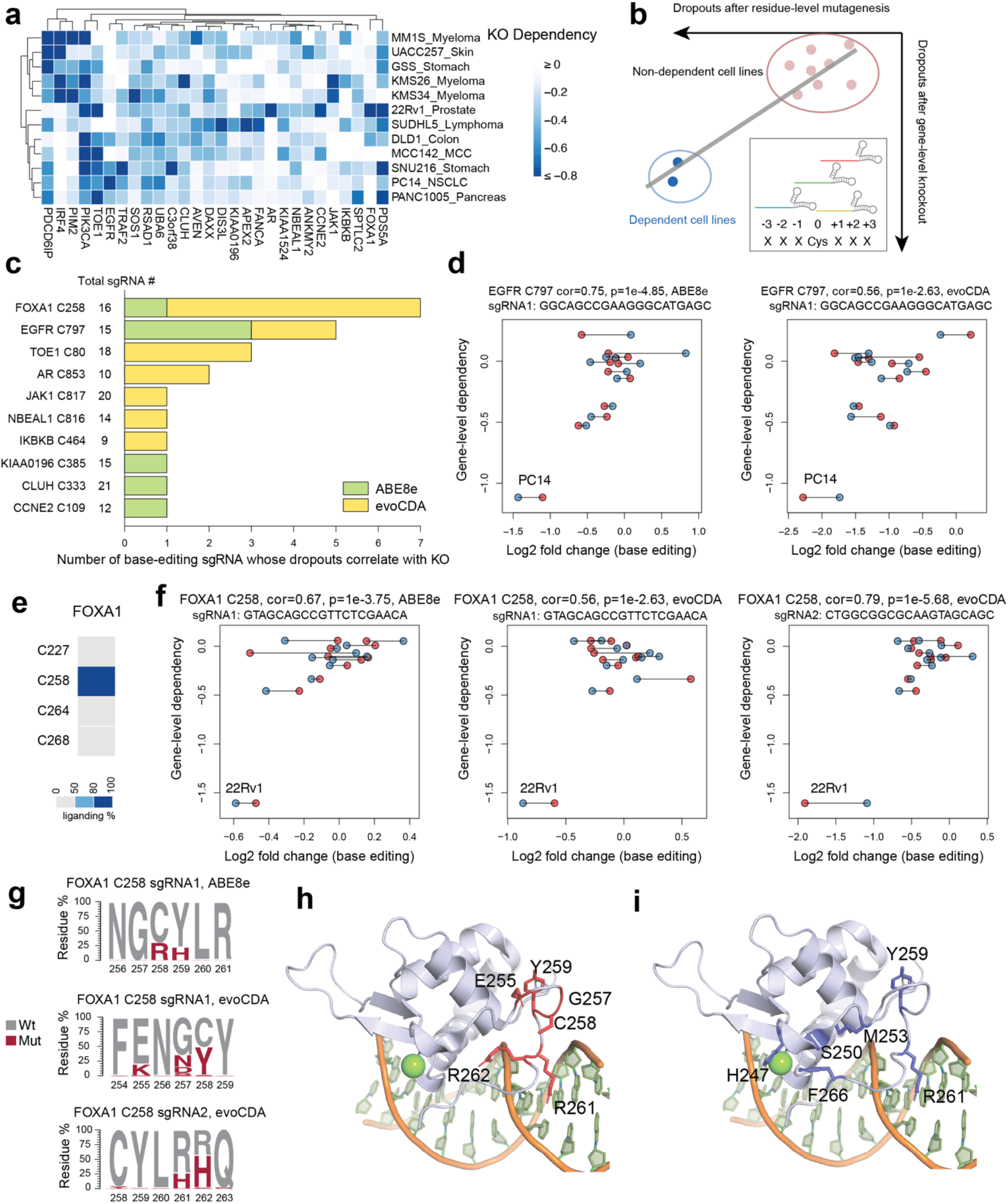
Base editing reveals essentiality of ligandable cysteine regions in Strongly Selective cancer dependency proteins. **a**, Heatmap showing Strongly Selective proteins with ligandable cysteines in the twelve-cell line panel. Strongly Selective designations are derived from the Cancer Dependency Map. **b**, Schematic showing how base-editing can assign essentiality to ligandable cysteine regions in Strongly Selective proteins. sgRNAs were designed to edit ligandable cysteines +/- 3 surrounding residues. An sgRNA is considered a hit if the resulting missense mutation leads to preferential dropouts of cancer cell lines that also show selective dependency on the corresponding protein containing the ligandable cysteine based on gene-level disruptions in the Cancer Dependency Map. **c**, Essential, ligandable cysteines identified in the twelve-cell line panel dropout screen, and the number of sgRNAs assigning essentiality to each cysteine. The total number of sgRNAs included in the screen for each cysteine region is also provided. **d**, Scatter plots for the EGFR_C797 region showing the correlation between base editing-induced cell line dropout data acquired herein and gene disruption-induced cell line dropout data in the Dependency Map. The two independent biological replicates for base-editing data in each of the twelve cell lines are shown in blue and red. Each plot shows a different sgRNA. **e**, The fragment electrophile ligandability profile for cysteines in FOXA1, showing ligandability of C258. **f**, Scatter plots for the FOXA1_C258 region showing the correlation between base editing-induced cell line dropout data acquired herein and gene disruption-induced cell line dropout data in the Dependency Map. The two independent biological replicates for base-editing data in each of the twelve cell lines are shown in blue and red. Each plot shows a different sgRNA. **g**, FOXA1 amino acid mutagenesis generated by different sgRNAs and the indicated base editors in the FOXA1-dependent cell line 22Rv1. The relative residue frequency five days after lentivirus infection was quantified using targeted genomic PCR and amplicon sequencing. Data represent average values of two independent experiments. **h**, **i**, FOXA1 DNA-binding domain homology model based on the crystal structure of FOXA3 in complex with DNA (PDB: 1VTN). The side chains of essential residues identified in the base-editing dropout screen and residues representing hotspot pro-cancerous mutations^38–41^ are highlighted in red and blue in **h** and **i**, respectively. Green ball represents Mg^2+^.

After 16 days of growth in the presence of sgRNAs, the cancer cell line pool was assessed for base-editing dropout events, and the magnitude of these effects was compared to reference dropout values for the corresponding gene disruptions contained in the Cancer Dependency Map. To account for different numbers of sgRNAs per cysteine region and avoid bias towards those with more sgRNAs, we performed multiple testing corrections by calculating false discovery rates (FDR) per site. We considered a cysteine to be essential when the base editing effect of an sgRNA targeting that cysteine’s region produced: i) a minimal LFC value of < −0.5 for the cell line(s) showing selective dependency on the corresponding gene product; and ii) the magnitude of base-editing dropout was greatest for the cell line(s) showing selective dependency on the corresponding gene product compared to other cell lines in the panel (Pearson correlation > 0.5 and associated FDR < 10%). Based on these criteria, we identified 10 ligandable cysteines displaying evidence of essentiality (**Fig 3c** and **Supplementary Table 3**). Among these essential, ligandable cysteines was EGFR_C797, which showed the expected preferential dropout in PC14 cells compared to other cell lines (**Fig. 3d**).

We noted that only a subset of the tested sgRNAs registered as hits for each essential, ligandable cysteine, and the total number of hit sgRNAs also varied widely for these residues (**Fig. 3c**). Consistent with differences in editing efficiency being a potential source for such variability, we found that, among 20 representative sgRNAs targeting the same cysteine regions that were individually cloned and arrayed, hit sgRNAs produced an average of 48% editing whereas non-hit sgRNAs produced only 13% editing (**Extended Data Fig. 3c**). These data indicated that the total number of hit sgRNAs per cysteine region may not correlate with the degree of essentiality, but rather the extent of editing efficiency achieved by the sgRNAs targeting this region, and, conversely, even a single hit sgRNA may be sufficient to assign essentiality to a cysteine region. Also supportive of this conclusion, we observed that JAK1_C817 was designated as essential by a single sgRNA (**Fig. 3c** and **Extended Data Fig. 3d, e**), and we have recently described covalent ligands targeting this cysteine that allosterically inhibit JAK1-dependent signaling in human cells^28^.

Among the other essential, ligandable cysteines was C258 of the pioneer transcription factor FOXA1 (**Fig. 3c, e, f**). FOXA1 plays a key role in the normal development of several endoderm-derived organs^43^ and represents a Strongly Selective dependency in subsets of prostate and breast cancer cell lines. Several base-editing sgRNAs registered as hits for FOXA1 (**Fig. 3c**), where preferential dropout was observed in the 22Rv1 prostate cancer cell line compared to other cell lines in the panel (**Fig. 3f**). We arrayed and quantified the genomeediting outcomes for three representative sgRNAs and found that they possessed distinct sets of missense mutations at C258 and/or neighboring residues (C258R+Y259H, E255K+G257N+C258Y, and R261H+R262H) (**Fig. 3g**). Structural analysis using a FOXA3 homology model^44^ indicated that C258, as well as the additional local residues altered by base editing, are part of the Wing2 region of the forkhead (FKHD) domain and reside in proximity to the FOXA1 DNA-binding site (**Fig. 3h**). Interestingly, the local region surrounding C258 is also enriched in oncogenic mutations that have been shown to remodel chromatin accessibility and gene expression, as well as affect the differentiation state, of breast and prostate cancer cells^45–48^ (**Fig. 3i**). Taken together, these data indicate that genetic alteration of the C258 region can modulate the activity of FOXA1, and the covalent ligandability of C258 further points to the possibility for small molecules to produce similar functional outcomes.

We were also intrigued by essential, ligandable cysteines discovered in Strongly Selective proteins that have less well-understood roles in cancer. One example was C80 in TOE1 (**Extended Data Fig. 3f**), a 3’ RNA exonuclease that regulates small nuclear RNA and telomerase RNA maturation^49,50^. Unlike other Strongly Selective proteins, such as FOXA1 or EGFR, which displayed essentiality in specific cancer lineages or in association with hotspot mutations, cell lines sensitive to TOE1 were distributed across various cancer types (https://depmap.org/portal/gene/TOE1?tab=dependency). In our twelve cell line panel, MCC142 (Merkel cell carcinoma) and PANC1005 (pancreatic adenocarcinoma) cells showed the greatest dependency on TOE1 as determined by gene-level dependency score, and our base-editing experiments produced a similar result for the TOE1_C80 region (**Extended Data Fig. 3g**). Because the codon for TOE1_C80 is on the edge of exon 4, we had limited options for sgRNAs targeting this residue without creating undesired splice-site mutations. Nonetheless, three sgRNAs targeting the C80 region registered as hits (**Fig. 3c**), and genomic sequencing revealed missense mutations that included C80Y and, to an even greater extent, E82K, E83K, and/or R84H (**Extended Data Fig. 3h**). Interestingly, the E82-E83-R84 sequence, but not C80, is conserved in the homologous nuclease PARN, despite this protein and TOE1 only sharing 30% overall identity (**Extended Data Fig. 3i**), and the crystal structure of PARN indicates these conserved residues are located distal from the nuclease active site (**Extended Data Fig. 3j**). Both TOE1 and PARN are parts of large macromolecular complexes required for RNA tail processing, and it is possible that the E82-E83-R84 stretch is important for protein-protein interactions or RNA substrate binding. The proximity of C80 – a paralog-restricted ligandable cysteine – to a stretch of highly conserved and genetically essential residues in the TOE1/PARN nuclease family suggests that covalent compounds targeting C80 have the potential to selectively impact the function of TOE1 over related nucleases.

### Quantitative reactivity profiling nominates essential cysteines with ligandability potential

In considering the scope and limitations of our approach, we grappled with the finding that such a small fraction of essential cysteines mapped by base-editing had evidence of covalent ligandability in chemical proteomic studies performed to date (109 ligandable, essential cysteines of > 1,350 total essential cysteines in the original global pooled screen). This can be explained, at least in part, by the limited diversity of chemistry evaluated in original ligandability maps, as chemical proteomic studies performed so far have only screened a handful of electrophilic fragments^10,16,17^ and therefore likely underestimate by a large margin the complete small-molecule interaction potential of cysteines in the human proteome. We also wondered, however, if other factors may be at play. For instance, some essential cysteines may have an intrinsically low likelihood of interacting with small molecules if, for instance, they serve structural roles at buried locations in proteins with limited or no access to solvent. Distinguishing such buried, essential cysteines from those that have greater potential to interact with small molecules would provide a way to prioritize cysteines for the future pursuit of chemical probes. With this goal in mind, we adapted established protocols for cysteine-directed ABPP^10,11^ to enable quantitative comparisons of cysteine reactivity with an iodoacetamide-desthiobiotin (IA-DTB) probe in denatured versus native proteomes (**Fig. 4a**). We hypothesized that this comparison may reveal bidirectional changes in cysteine reactivity that were informative for classifying different types of functional residues: i) essential cysteines showing increased reactivity following protein denaturation may serve structural roles, but be inaccessible to solvent in the native protein state; and ii) essential cysteines showing decreased reactivity following protein denaturation may be solvent-accessible residues proximal to pockets that promote enhanced IA-DTB reactivity.

**Fig. 4.**
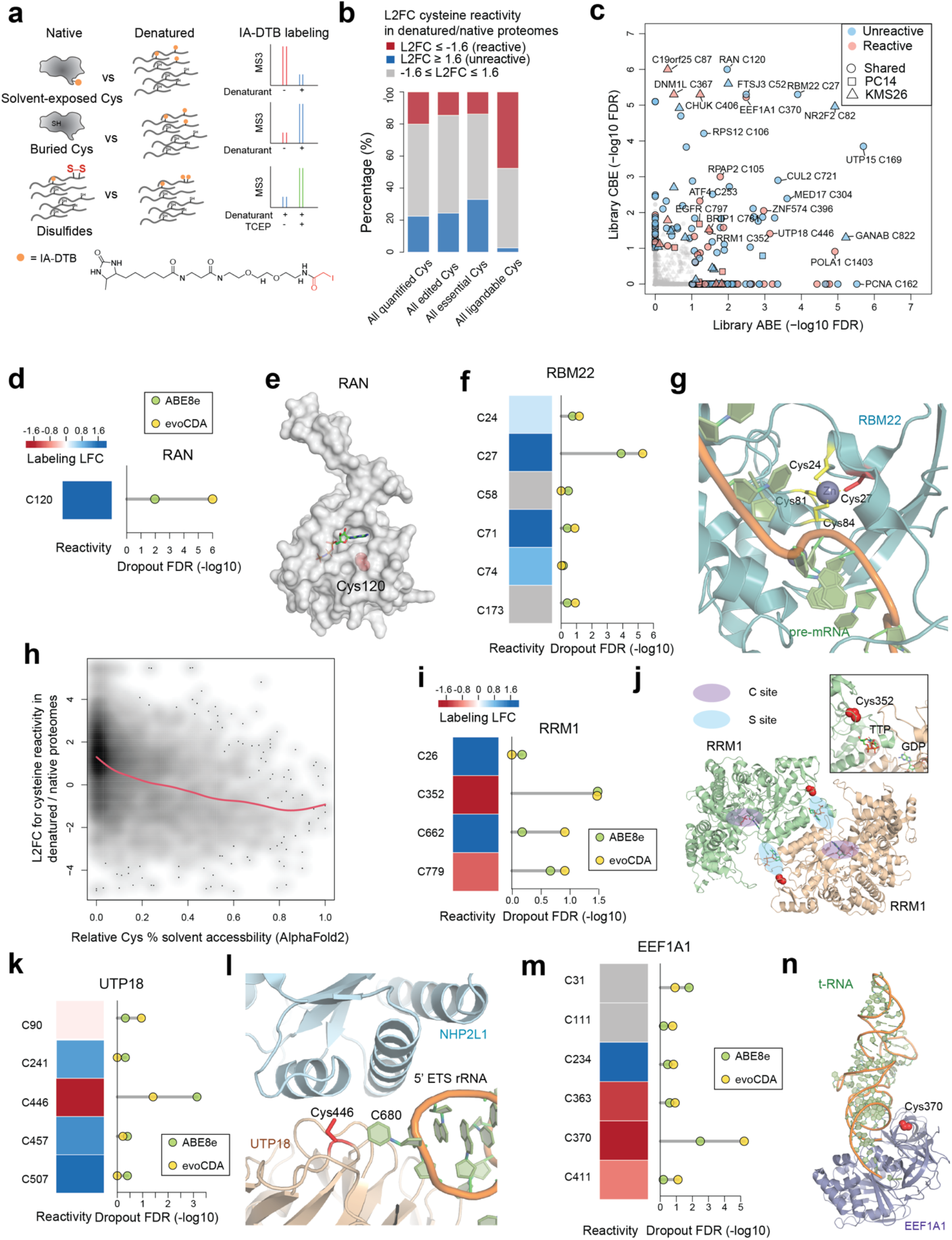
Prioritizing essential cysteines with ligandability potential by quantitative cysteine reactivity profiling in native and denatured proteomes. **a**, Schematic showing an ABPP workflow to distinguish solvent-exposed and buried cysteines based on quantitative differences in IA-DTB reactivity in native and denatured proteomes. **b**, Stack bar plots summarizing the distribution of reactivity changes for the indicated categories of cysteines, including 1) all cysteines quantified in the ABPP experiments of native versus denatured proteomes; 2) all cysteines that were base-edited; 3) all essential cysteines (i.e., cysteines with editing-induced dropouts of LFC<-0.6, FDR<10%); and 4) all ligandable cysteines with electrophilic fragments. **c**, Summary of editing-induced dropout values for cysteines showing substantially greater or lesser reactivity in native proteomes (designated as reactive and unreactive, respectively). Substantial reactivity changes were defined as cysteines showing denaturing/native L2FC scores < −1.6 (reactive) or > 1.6 (unreactive). Cysteine-directed ABPP data represent average values from at least three independent biological experiments, and the cysteine essentiality data were derived from the large-scale base editing screen described in **Fig 2**. **d**, The reactivity in denatured/native proteomes (left) and significance of dropout (right) for the edited C120 in RAN. **e**, Structure of RAN in complex with GNP (GTP analogue) (PDB: 5JLJ). The side chain of essential, unreactive (buried) cysteine C120 is highlighted in red. **f**, The reactivity in denatured/native proteomes (left) and significance of dropout (right) for cysteines in RBM22. **g**, Structure of RBM22 in complex with pre-mRNA as part of the human spliceosome (PDB: 5XJC). The side chain of identified essential, unreactive (buried) C27 is highlighted in red. **h**, Density plot showing the association between predicted cysteine solvent accessibility based on AlphaFold2 protein structures (threshold of residue confidence pLDDT > 70) and quantified cysteine reactivity values (denatured/native proteomes) from ABPP experiments. A fitted smoothing spline is shown in red. **i**, The reactivity in denatured/native proteomes (left) and significance of dropout (right) for cysteines in RRM1. **j**, Structure of RRM1 bound to effector metabolite TTP at the specificity site (S site) and substrate GDP at the catalytic site (C site) (PDB: 3HND). The side chain of essential, reactive (solvent-exposed) cysteine C352 is highlighted in red. **k**, The reactivity in denatured/native proteomes (left) and significance of dropout (right) for cysteines in UTP18. **l**, Structure of UTP18 as part of the human small subunit processome (PDB: 7MQ8). The side chain of essential, reactive (solvent-exposed) cysteine C446 is highlighted in red. **m**, The reactivity in denatured/native proteomes (left) and significance of dropout (right) for cysteines in EEF1A1. **n**, Structure of EEF1A1 in complex with tRNA as part of the human ribosome complex (PDB: 6ZMO). The side chain of essential, reactive (solvent-exposed) cysteine C370 is highlighted in red.

Denaturation of the KMS26 and PC14 proteomes was performed with sodium dodecyl sulfate (SDS) or urea, and cysteine reactivity profiles were compared to native cell proteomes by multiplexed quantitative (tandem mass tagging (TMT)) ABPP. An initial evaluation of SDS- and urea-treated proteomes revealed that each denaturant induced similar cysteine reactivity changes (**Extended Data Fig. 4a**), and we therefore combined data from both denaturation conditions for a more detailed comparison to native proteomes (**Supplementary Table 4**). Across ~29,000 quantified cysteines, ~20% and 22% showed substantial (> 1.6-log2-fold change (LFC)) decreases or increases in IA-DTB reactivity in denatured proteomes, respectively (referred to hereafter as reactive and unreactive cysteines, respectively; **Fig. 4b** and **Supplementary Table 4**). A similar distribution of cysteine reactivity changes was observed for the ~5,000 quantified cysteines that had been edited in our dropout screens (**Fig. 4b** and **Supplementary Table 4**). In contrast, we found that a greater relative fraction of the ~500 essential cysteines quantified by ABPP were unreactive (**Fig. 4b** and **Supplementary Table 4**), indicating an enrichment in cysteines that were preferentially accessible to the IA-DTB probe in unfolded protein states. Consistent with previous studies^10^, a striking converse relationship was observed for the ~5,500 ligandable cysteines quantified by ABPP, which were strongly enriched in reactive cysteines with virtually no representation of unreactive cysteines (**Fig. 4b** and **Supplementary Table 4**).

Structural analysis of representative proteins with essential, unreactive cysteines (blue proteins, **Fig. 4c**) supported the premise that these residues are buried with limited access to solvent. For instance, an essential cysteine C120 in the GTP-binding nucleocytoplasmic transport protein RAN showed a seven-fold increase in IA-DTB reactivity after denaturation and has a solvent-accessible surface area in the folded protein structure of 0 Å^2^ (**Fig. 4d, e**). Likewise, the essential cysteine C169 in the ribosome biogenesis factor UTP15, which also showed a striking denaturation-induced increase in reactivity, has its accessibility to solvent restricted by interactions with the associated proteins NOC4L and RPS18^51^ (**Extended Data Fig. 4b, c**). Additional essential, unreactive cysteines were found to chelate metals, such as C27 of the splicing factor RBM22, which is part of a zinc finger that interacts with pre-mRNA substrates^52^ (**Fig. 4f, g**). Finally, using AlphaFold2-generated structural models^53,54^, specifically residues that passed a threshold of confidence (pLDDT > 70), we predicted the solvent accessibility for > 10,000 cysteines quantified in our ABPP experiments (**Supplementary Table 4**), which revealed that unreactive cysteines were generally less accessible to solvent (**Fig. 4h**, Pearson correlation = −0.38, p < 2.2e^-16^). These data, taken together, indicate that a substantial number of essential cysteines may lack evidence of ligandability because they serve structural roles at buried sites within proteins that have limited accessibility to small molecules.

Our analysis conversely revealed a set of 68 essential, reactive cysteines (red proteins, **Fig. 4b, c**; also see **Supplementary Table 4**). Considering that heightened reactivity in native proteomes was a feature strongly enriched in ligandable cysteines (**Fig. 4b**), we consider essential cysteines showing this property as prioritized targets for the future pursuit of covalent probes. An initial analysis of these essential, reactive cysteines revealed several located at macromolecular interfaces. The essential, reactive cysteine C352 in ribonucleotide reductase 1 (RRM1) (**Fig. 4c, i**), for instance, resides at the dimerization interface of this enzyme in proximity to an allosteric regulatory site where specificity effector nucleoside triphosphates bind to regulate RRM1 activity^55^ (**Fig. 4j**). Other essential, reactive cysteines were located at protein-RNA/DNA interfaces, including C446 in UTP18 (**Fig. 4c, k**), a component of human small subunit processome. C446 resides at the UTP18 and NHP2L1 interface in complex with a 5’ external transcribed spacer (ETS) RNA that structurally facilitates initial stages of processome assembly^51^ (**Fig. 4l**). Similarly, the essential, reactive cysteine C370 in EEF1A1 (**Fig. 4c, m**), a protein that promotes GTP-dependent binding of aminoacyl tRNA to ribosomes^56^, was located at the EEF1A1-tRNA interface (**Fig. 4n**). Essential, reactive cysteines were also identified in Strongly Selective cancer dependencies, such as C761 in BRIP1 (**Fig. 4c**, **Extended Data Fig. 4d**), a DNA helicase that is required for DNA double-stranded break repair and the maintenance of chromosomal stability^57^. Based on a homology model generated from the related protein ERCC2, we localized C761 to the helicase domain at a site predicted to be proximal to BRIP1-DNA substrate interactions (**Extended Data Fig. 4e, f**).

Together, these data show how the integration of base editing data with global maps of cysteine reactivity in native and denatured proteomes, can illuminate solvent-accessible, essential cysteines as prioritized targets for future chemical probe development.

## Discussion

ABPP and related chemical proteomic methods have discovered a wide array of cysteine residues in diverse protein classes that can be targeted by electrophilic small molecules^10,16–18,58^. It remains challenging, however, to determine the functional consequences of such covalent liganding events, especially for proteins lacking established biochemical or cellular assays. Additionally, initial hit ligands discovered by chemical proteomics often represent simple electrophilic fragments that lack sufficient potency and selectivity for immediate use in cell-based functional assays. Here, we describe a genome editing approach for globally assessing the functionality of ligandable cysteines in proteins required for cancer cell growth. Key to the implementation of our strategy was the knowledge afforded by - i) ABPP of site-level resolution of covalent compound-cysteine interactions across the human proteome^10,16–18,58^; and ii) the Cancer Dependency Map of protein-encoding genes required for cancer cell proliferation^5^ - which, together, enabled focused, base editing-mediated mutational screens of ligandable cysteines and surrounding regions to assess their essentiality for cell growth.

We chose base editing to install mutations because this technology can introduce nucleotide changes for residue-level functional assessment, while minimizing DNA-double-strand breaks^30,31^ and uncontrolled mixture of indels^59^ that are observed with homology-directed repair (HDR) CRISPR-Cas9 nuclease systems. Indeed, base editing has been used to evaluate cancer-associated missense mutations emerging from clinical genomics data^60–62^, as well as to discover missense mutations that cause resistance to targeted therapeutics^62^. We were also fortunate that the ABE and CBE systems produced non-conservative mutations of cysteines (Cys-to-Arg or Cys-to-Tyr) that were anticipated to have good potential to alter the functions of proteins. Recently, an alternative HDR-based oligo recombineering approach was described to assess the functionality of cysteines in *Toxoplasma gondii*^63^. This approach is well-suited for focused, array-based screens, particularly in haploid systems, but currently lacks the throughput and efficiency to perform large-scale analyses of thousands of cysteines in diploid mammalian cells.

Our base editing study assigned essentiality to >110 ligandable cysteines, including those targeted by covalent drugs that have been clinically approved for treating cancer (e.g., EGFR_C797 and XPO1_C528). The discovery of essential, ligandable cysteines in proteins encoded by genes with Strongly Selective designations in the Cancer Dependency Map, such as CHUK, FOXA1, and TOE1, is particularly intriguing, as the development of more advanced covalent probes for these proteins would not only help to define their specialized pro-tumorigenic functions, but also may serve as starting points for targeted cancer therapies. Considering further that a subset of these ligandable cysteines (e.g., C258 in FOXA1^45–47^) are proximal to hotspot mutations in cancer, we wonder whether it might be possible to create covalent probes that specifically target such pro-tumorigenic mutant forms of proteins, as has been done for EGFR^24^ and KRAS^26,64^. This could be important for translational pursuits of FOXA1, as C258 is conserved in the paralogous proteins FOXA2 and FOXA3.

The base editing of more than 1,100 ligandable cysteines in cancer dependency proteins did not affect cancer cell growth in our screens. One interpretation of these results is that ligands targeting these cysteines may not substantially affect protein function. However, multiple technical caveats should also be considered. First, some covalent ligand-cysteine interactions may exert functional effects that qualitatively or quantitatively differ from those produced by genetic mutation of proteins. Indeed, our base-editing strategy was limited to converting cysteines to arginine or tyrosine, and these mutations may not fully mimic the functional effects of engagement of cysteines by covalent small molecules, which can create much more chemically diverse modifications. We attempted to account for this difference, at least in part, in our focused screen of ~30 ligandable cysteines in Strongly Selective proteins by using base editors with expanded targeting windows, allowing for mutation of additional residues that neighbor ligandable cysteines, which we anticipated would more fully assess the functionality of the local cysteine region (as we found to be the case for EGFR_C797 and XPO1_C528; **Fig. 1e, g**). Additionally, some ligandable cysteines that were designated as non-essential by base-editing screens may lack interpretability if insufficient gene editing of that cysteine region was technically achieved in our screens. Future genetic interrogation of cysteine essentiality could benefit from editors with improved efficiency that use, for instance, PAM-less Cas9 and methods that provide direct monitoring of editing efficiencies in parallel^60,61^.

Our base editing screens also identified a large number of essential cysteines (~1,250) for which ligands have not yet been discovered. While many of these cysteines can continue to be evaluated by chemical proteomic methods for interactions with distinct types of electrophilic compounds, this process is admittedly low-throughput and somewhat serendipitous in outcome. On the other hand, determining which essential, but not yet liganded, cysteines have the highest potential for targeting by covalent chemistry could enable more focused screens compatible with evaluating larger compound libraries. We have addressed this goal by globally quantifying the relative reactivity of cysteines with the IA-DTB probe in native versus denatured states, which we found to provide a robust way to prioritize essential cysteines with greater ligandability potential. We specifically discovered that heightened reactivity following denaturation was correlated with cysteines that are buried (i.e., solvent-inaccessible) in protein structures, and, conversely, cysteines with established ligandability were much more likely to show impaired reactivity after denaturation. Perhaps most striking was the near absence of ligandable cysteines showing heightened reactivity following denaturation. Accordingly, we proposed that essential cysteines displaying this property are much less likely to be targetable by covalent chemistry, at least as pertains to the major proteoforms harboring these cysteines in the cancer cells investigated herein. We also speculate that denaturation-induced increases in cysteine reactivity may more generally flag a category of residues for which essentiality reflects mutation-induced disruption of protein folding and/or expression.

In summary, by integrating base editing and chemical proteomic technologies, we have generated a rich resource of cysteines and cysteine regions that are essential for the growth of cancer cells alongside a status report on the current and future potential for targeting these cysteines with covalent small molecules. From a translational perspective, essential, ligandable cysteines in proteins with a Strongly Selective designation in the Cancer Dependency Map are particularly interesting, as chemical probes targeting these proteins would be expected to preferentially affect specific subtypes of cancer as opposed to acting as general disruptors of cell growth. We should note, in this regard, that our twelve-cell line panel only captured a limited subset of the total number of Strongly Selective genes assigned by the Cancer Dependency Map, and the future base editing and chemical proteomic analysis of additional cancer cell lines is likely to uncover many other essential, ligandable cysteines in proteins that preferentially regulate the growth of specific cancers. Even essential, ligandable cysteines identified in Common Essential proteins may have translational relevance, if, for instance, partial blockade of protein function by small molecules can produce a more selective effect on cancer growth than total gene/protein disruption^65^. For instance, gene-level loss of RRM1 or EEF1A1 – two Common Essential proteins with essential, ligandable cysteines (C352 and C370, respectively) – shows quantitatively greater growth effects in cancer cell lines with reductions in copy number (RRM1; **Extended Data Fig. 4g**) or transcriptional expression of a paralog (EEF1A2; **Extended Data Fig. 4h**), pointing to molecular markers that may constitute synthetic lethality relationships. In the future, we also envision performing integrated base-editing and chemical proteomic screens coupled to high-throughput readouts of other features of cell biology beyond proliferation and, through doing so, further enriching our understanding of functional cysteines that can be targeted by covalent chemistry.

## Data availability

The raw sequencing data have been deposited to NCBI GEO. The raw proteomics data have been deposited to PRIDE.

## Code availability

Custom code used in the analysis will be made available on github prior to publication.

## Acknowledgements

We thank Dr. John Doench (Broad institute) and Dr. Ji Luo (NIH) for helpful discussions regarding CRISPR library cloning. We also thank Dr. Bruno Melillo and Dr. Kristen DeMeester for feedback related to electrophile probes used in the study. This work was supported by the NIH (R35 CA CA231991, U01 AI142756, RM1 HG009490, R35 GM118062), the Damon Runyon Cancer Research Foundation (DRG: 2406-20), and the Howard Hughes Medical Institute (HHMI).

## Author Contributions

H.L. and B.F.C. conceived the study and wrote the manuscript. H.L., J.R.R, S.J.W performed the experiments. H.L., K.T.Z., T.P.H. and D.R.L. contributed in the design and interpretation of base editing experiments. H.L., G.M.S. and B.L. contributed to data analysis. All authors edited and approved the manuscript. B.F.C. supervised the study.

## Methods

### Cell culture

Authenticated human cancer cell lines were collected from ATCC, ECACC and JCRB cell banks. All cancer cell lines were grown in RPMI-1640 (Gibco) media using standard cell culture conditions (37°C, 5% CO2) and were free of microbial contamination including mycoplasma. All media were supplemented with 100 U/ml penicillin, 100 μg/ml streptomycin (Gibco), 10% FBS (Omega Scientific), and 2 mM GlutaMAX (Gibco).

### Lentivirus particle production

The plasmids including all-in-one base editor vector, lentiviral packaging vector (pCMV-dR8.91) and envelope vector (VSV-G) were mixed at 9:6:1 ratio in OPTI-MEM media. 3ul of 1mg/ml PEI-MAX transfection reagent (Polysciences) was added per ug of total plasmids. After 20 min of incubation at room temperature, the mixture was dripped gently to 293T cells at 50% confluence. After 8 hrs, the media was changed to fresh DMEM media with 30% FBS plus penstrep and 2mM GlutaMAX. The virus was collected both 2 days and 3 days later and 0.45 μm syringe filters (Millipore) were used to eliminate cells. The virus-containing supernatant was then aliquoted and frozen at −80°C before use.

### Arrayed base editing

On day 0, cells were seeded in a 96-well plate at a density of 0.5-1e4/well and were mixed with virus supernatant with a final polybrene concentration at 8 μg/ml. Spin infection was performed by centrifugation at 900g for 1 hour at 30C. On day 1, the virus containing media was removed, and fresh media was added. On day 2, puromycin was added to start the selection. The transduced cells were collected on day 5 after PBS wash and were stored at −80°C before use.

### Targeted genomic sequencing

Genomic sites with base editing were amplified and indexed using a two-step PCR method as previously described with minor modifications^66^. Briefly, 100ul lysis buffer (10 mM Tris pH 7.5, 0.5% Tween 20, 0.02% SDS plus 20 ug/ml freshly added proteinase K) was added per 1e5 cells and the samples were incubated at 55°C for 2 hr before heat inactivation at 95°C for 30 min. Primers with illumina adapters were used for PCR1. Specifically, in each reaction, 5ul of genomic DNA extract, 12.5 ul Phusion PCR master mix (ThermoFisher), 1.25ul of 10uM forward and 1.25ul of 10uM reverse primers were added to a total volume of 25ul. The PCR1 reactions were carried out as following: 95°C for 3 min, repeated cycles of (20s at 95°C, 20s at 60°C, 25s at 72°C), followed by final extension at 72°C for 2 min. PCR1 products were cleaned using Ampure beads (Beckman) according to manufacturer’s instructions and were eluted in 50ul of 10mM Tris (pH=7.5). For each PCR2 reaction, 5 ul of PCR1 product, 12.5 ul Phusion PCR master mix (Thermo Scientific), 1.25ul of 10uM forward and 1.25ul of 10uM reverse index primers were added to a total volume of 25ul. The PCR2 reactions were carried out as following: 98°C for 3 min, then repeated cycles of (20s at 95°C, 20s at 60°C, 25s at 72°C), followed by final extension at 72°C for 2 min. The PCR2 products were then pooled and cleaned using Ampure beads. The library was quantified using PicoGreen dsDNA assay kits (ThermoFisher) and sequenced on an Illumina Miniseq instrument with 10% PhiX spike-in. Paired-end reads were demultiplexed based on combinatorial dual indexes in Local Run Manager (Illumina). The genome editing quantification was performed using

CRISPResso2^67^ (https://crispresso.pinellolab.partners.org). The parameters were set as following: CRISPResso --fastq_r1 {r1_file} --fastq_r2 {r2_file} --amplicon_seq {amplicon_seq} −g {guide} −wc −14 −w 20 −q 30 --base_editor_output

### Design of pooled base editing libraries

The dependency “CERES” scores were downloaded from the Cancer Dependency Map (21Q3, https://depmap.org/portal/). To design the libraries in this study, we focused on protein-coding transcripts annotated by the GENCODE database (http://www.gencodegenes.org/). For each protein, we selected the principal isoform based on the APPRIS database^68^. We further defined the approximate editing window center to be 15 nucleotides upstream of the NG PAM with 5nt width for ABE8e and 10nt width for evoCDA. The sgRNA sequences were selected if the editing window were predicted to create missense mutations to the target position or nearby residues if relevant. We then appended BsmBI sites to allow restriction digestion and PCR primer binding sites to allow oligo pool amplification. The final oligo structure is: 5’-(primer forward)CGTCTCACACCG(sgRNA, 20 nt)GTTTCGAGACG (primer reverse).

### Cloning of pooled base editing libraries

All-in-one lenti-ABE8e-NG and lenti-evoCDA-NG vectors with sgRNA inserts were made with golden gate cloning. Briefly, the plasmids were digested with BsmBI (NEB) following manufacturer’s recommendations and the backbone DNA with sticky ends were then separated in 1% agarose gel and purified for later use. The sgRNA oligo pool with restriction sites and flanking primer sequences was synthesized by Twist Bioscience and was amplified using Q5 polymerase (NEB). The PCR reaction was carried out as following: 98°C for 3 min, then repeated cycles of (20s at 95°C, 20s at 53°C, 20s at 72°C), followed by final extension at 72°C for 2 min. The PCR products were cleaned using DNA Clean & Concentrator (Zymo). The ligation product was assembled as following: 5 ng insert, 1 ug digested backbone, 1x Tango buffer, 1mM DTT, 1mM ATP, 1ul Esp3I (ThermoFisher), 1ul T7 ligase (Qiagen Beverly) in a total volume of 50ul. The ligation products were cleaned using isopropanol precipitation and were electroporated into electrocompetent cells. After 1 hr incubation in recover media at 37°C, the bacteria cells were spread onto 145mm * 145mm plates for overnight culture at 30°C. For each sgRNA, at least 1,000 colonies were obtained to get sufficient library representation. After 20hr, the bacteria cells were scraped and centrifuged at 3,000g for 10 min and the plasmids were then extracted (Qiagen). Cloning of individual sgRNA base editor construct was based on similar procedure with reduced reagent input.

### Cysteine reactivity profiling of native versus denatured proteomes evaluated by ABPP

10 million cells (PC14 or KMS26) were collected for each condition or replicate after three PBS washes. Each frozen cell pellet was mixed with 300 ul PBS and was sonicated with 3*8 pulses on ice. The protein was then normalized to 1 mg in a total of 500 ul volume with different treatments (8M urea at 65°C for 15 min, 1% SDS at 95 °C for 5 min, or 1% SDS+2 mM TCEP at 95°C for 5min). The native groups were mock treated with PBS and were kept on ice before further use. After equilibrating to room temperature, the samples were then treated with 5 ul 10 mM stock of IA-DTB (SCBT) and were incubated at room temperature for 1 hr. To precipitate proteins, 500ul cold methanol and 200ul cold chloroform was added to each tube. The samples were vortexed and then centrifuged at 16,000g for 30 min at 4°C. After removing the liquid phase, the protein disk was washed with cold methanol and was centrifuged again at 16,000g for 30 min at 4°C. The liquid phase was then aspirated, and the pellet is frozen at −80°C for later use. The subsequent sample processing and LC-MS instrumentation is the same as the previously described protocol for cysteine ligandability profiling^17,29^. Briefly, the protein pellets were reduced by DTT, alkylated by iodoacetamide, and digested by trypsin overnight. The peptides were then enriched using streptavidin, labeled with TMT tags, followed by desalting using Sep-Pak C18 cartridges. The peptides were fractionated using high-pH HPLC methods and were then analyzed in Orbitrap Fusion™ Mass Spectrometer (ThermoFisher).

### Analysis of cysteine reactivity profiling data

The MS2 and MS3 files were extracted from the raw data files using RAW Converter (version 1.1.0.22; available at http://fields.scripps.edu/rawconv/) and were uploaded to Integrated Proteomics Pipeline (IP2). The data files were then processed using the ProLuCID program based on a reverse concatenated, non-redundant version of the Human UniProt database (release 2016-07). Cysteine residues were searched with a static modification for carboxyamidomethylation (+57.02146 Da). N-termini and lysine residues were searched with a static modification corresponding to the TMT tag (+229.1629 Da). To search for the cysteine IA-DTB labeling, a dynamic modification (+398.25292 Da) was used. The census output files from IP2 were further processed by aggregating TMT reporter ion intensities to obtain signals based on unique peptides that are further annotated with protein-cysteine residue numbers. The resulting data were then median normalized per TMT channel and log2 fold changes between the native versus denatured conditions were calculated for each cysteine. P values associated with quantified cysteines were obtained using two-sided Student’s t-tests and FDR values were calculated using the Benjamini-Hochberg procedure. Those cysteines with FDR>10% were excluded from further analysis. The denaturing/native L2FC summary scores were calculated per cysteine using available datasets based on both model cell lines (PC14 and KMS26) and different denaturants (SDS and urea). Substantial reactivity changes were defined as cysteines showing denaturing/native L2FC scores < −1.6 (reactive) or > 1.6 (unreactive).

### Analysis of cysteine solvent accessibility

The cysteine solvent accessibility data was extracted based on the predicted models in the AlphaFold Protein Structure Database (https://alphafold.ebi.ac.uk/). The residue-level solvent accessibility scores were calculated using DSSP from BioPython (https://biopython.org/). Only cysteine residues with confidence scores pLDDT > 70 were used in the analysis. A smoothing spline was fitted using the smooth.spline function in R with cross-validations (lambda=0.01).

### Cysteine ligandability with electrophilic fragments evaluated by ABPP

The list of ligandable cysteines derived from previous reactive fragment electrophile profiling studies^10,16,17^ was used without change of analysis workflow or threshold. The additional cysteine-directed ABPP experiments performed in this study were the based on the previously described protocol^17^. These data were generated by treating three different model cancer cell lines (RAMOS, 22Rv1 and MCF7) with covalent ligands including 200uM KB02 and KB05^10^ *in situ* for 1 hour. The unliganded cysteine were then labeled by iodoacetamide desthiobiotin (IA-DTB), following by streptavidin enrichment, trypsin digestion and multiplexed proteomic quantification. We define a site to be ligandable if the fragment can engage > 50% of the IA-DTB-reactive cysteine in at least one of the model cell lines.

### Screens of base-editing libraries

On day 0, cells were seeded in 12-well plates at a density of 0.5-1e6/well and were infected in virus supernatant containing 8 μg/ml of polybrene. Spin infection was performed by centrifugation at 900g for 1 hour. One day after infection, 20% of cells were frozen as pellets for library normalization and the rest of cells were split to 15cm plates or T175 flasks. On day 2, puromycin was added and was maintained throughout the entire screen. About 30-50% infection rate was achieved in the screen to get an optimal multiplicity of infection (MOI). Each sgRNA was screened in at least 1000 cells and the cells were cultured for additional 14 days. Genomic DNA from the cell pellets was purified using the NucleoSpin blood L kits (Macherey-Nagel) according to the manufacturer’s instructions and was quantified using PicoGreen dsDNA assay kits (ThermoFisher).

The number of total PCR reactions performed for each cell line is calculated by sgRNA number in the library * 1000 cells /1e6 and was rounded to the next integer. For each targeted sgRNA cassette amplification PCR, 5ug of genomic DNA, 50ul Phusion PCR master mix (ThermoFisher), 5ul of 10uM forward and 5ul of 10uM reverse primers with illumina adaptors and indexes were added to a total volume of 100ul. The PCR reactions were carried out as following: 98°C for 3 min, repeated cycles of (20s at 98°C, 20s at 53°C, 20s at 72°C), followed by final extension at 72°C for 10 min. The PCR products were then pooled and cleaned using Ampure beads (Backman) and were sequenced in an Illumina sequencer (HiSeq, NextSeq or NovaSeq) with 20% PhiX spike-in.

### Analysis of pooled screen data

The frequency of sgRNA on day 1 and day 16 were counted using PoolQ (Broad Institute, https://portals.broadinstitute.org/gpp/public/software/poolq) and were normalized by total reads per sample to get counts per million (CPM). The statistical significance of cysteine dropouts was calculated by comparing day 16 vs day 1 average log2 fold CPM changes of targeting sgRNA versus non-targeting sgRNA (null distribution). Here, the targeting sgRNAs include those that are predicted to mutate the cysteine of interest or nearby residues. Additionally, the sgRNAs that are predicted to introduce stop codons or make mutations near splicing sites were excluded from further analysis. The log2 fold CPM changes of non-targeting sgRNA were randomly sampled to get a null distribution based on the number of targeting sgRNA. The p value for each site was then estimated as the percentage of null observations that have greater average dropouts than the observed average dropout. The false discovery rate (FDR) was calculated using the Benjamini-Hochberg procedure for all edited cysteines in each protein.

For the cell panel screens, the sgRNA data was similarly counted and normalized. The day 16 vs day 1 dropouts were calculated for each sgRNA in each cell line. To select hit sgRNAs for each dependency, the base editing-induced dropout log2 fold CPM changes in the evaluated cell lines should correlate with the corresponding gene knockout-induced dropout scores (CERES 21Q3 release, https://depmap.org/portal/) with Pearson correlation greater than 0.5 and FDR < 10%.

---

**Extended Data Fig. 1.**
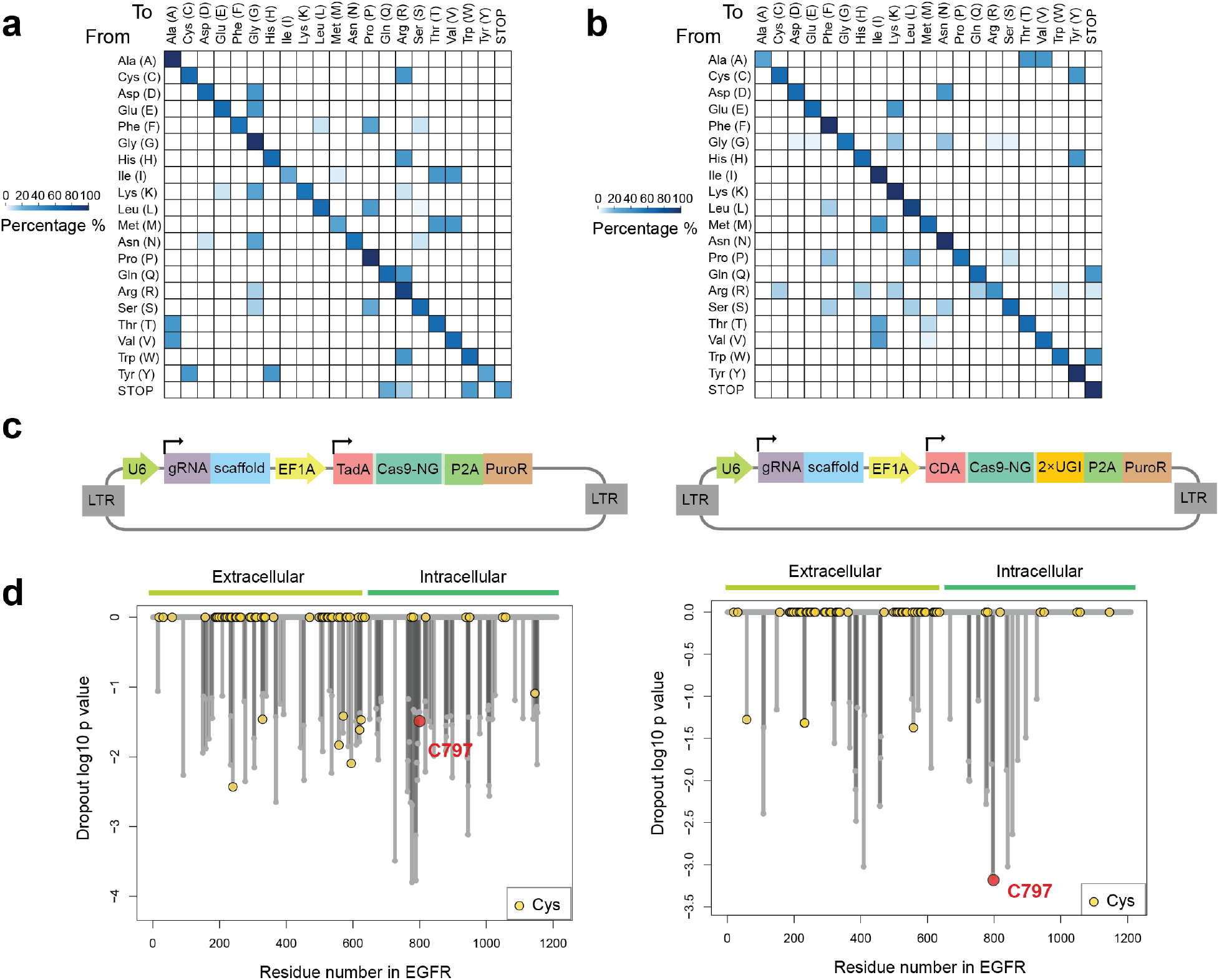
Base editing assigns essentiality to cysteines targeted by anticancer drugs. **a**, **b**, Heatmap showing the mutagenesis effects of Adenine Base Editors (ABE) (**a**) and Cytosine Base Editors (CBE) (**b**) on different amino acid residues. The total percentage is normalized to 100% per row. **c**, Schematic showing the design of lentivirus vectors delivering base editor libraries used in this study. CDA: cytidine deaminase; TadA: deoxyadenosine deaminases; UGI: uracil glycosylase inhibitor; PuroR: puromycin resistance gene. **d**, Waterfall plots for ABE (left) and CBE (right) libraries showing the dropout significance calculated per EGFR residue between day 16 vs day 1 from saturated scanning experiments. C797 is marked as a red circle, and other cysteine residues are highlighted as yellow circles. Data represent average values of two independent experiments. P values were calculated using Student’s t test.

**Extended Data Fig. 2.**
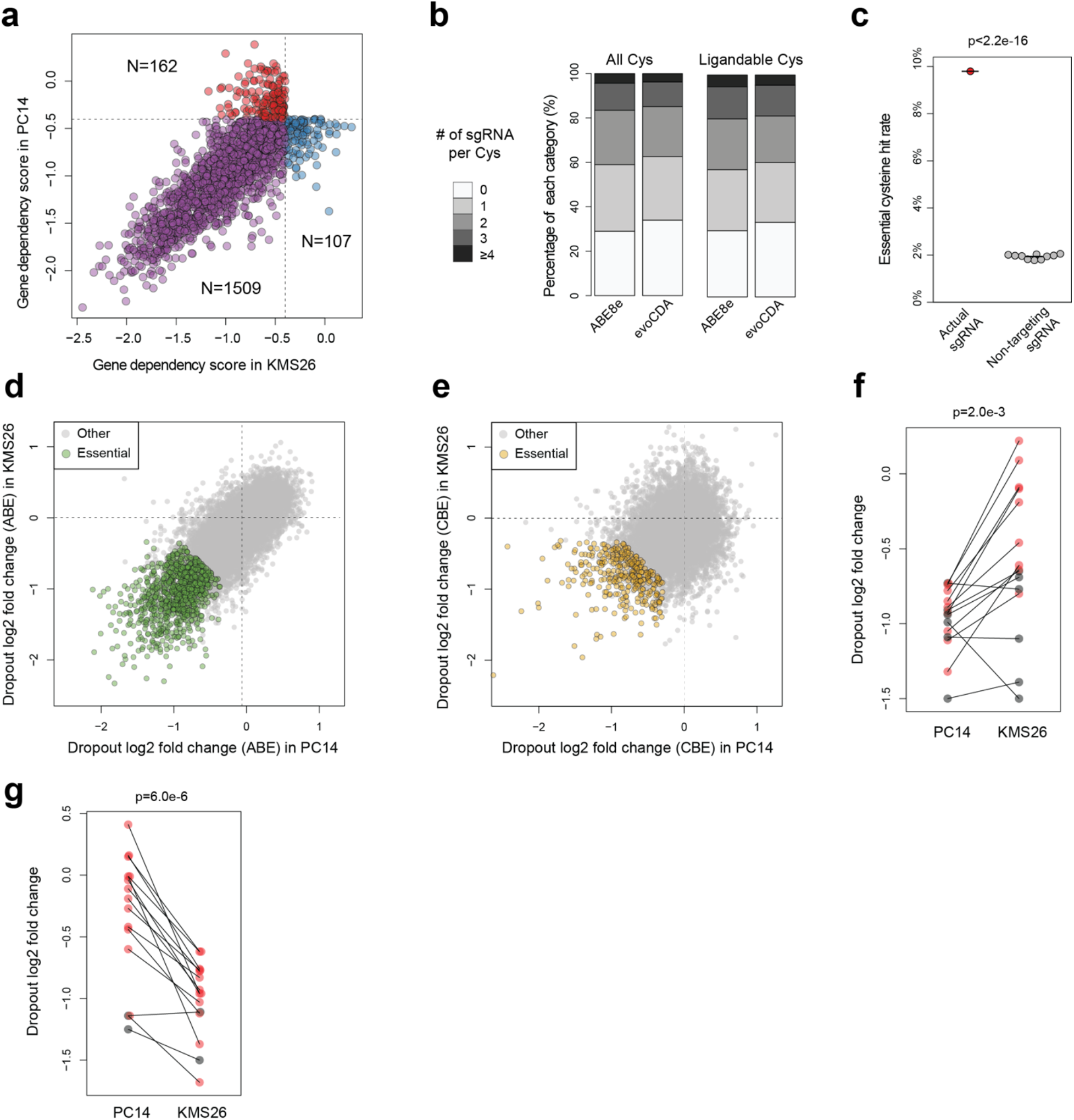
Global discovery of essential cysteines in cancer dependency proteins. **a**, Scatter plot showing the dependency of PC14 and KMS26 cells on genes included in the base-editing dropout screens investigating cysteine essentiality. Each gene is represented as a point (data source: the Cancer Dependency Map; https://depmap.org/portal/, 21Q3 release). Only genes with CERES dependency scores <-0.4 in at least one of the two cancer cell lines were used in the analysis. **b**, Stacked bar plots summarizing the number of designed sgRNAs per cysteine. (Left) All cysteines in cancer dependency proteins; (right) ligandable cysteines in cancer dependency proteins. **c**, Beeswarm plot comparing the essential cysteine hit rate (cutoff: LFC <= −0.6 and FDR < 10%) using acquired sgRNA data targeting cysteines in cancer dependency proteins or non-targeting sgRNA data randomly sampled from the ABE or CBE library. The analysis was performed 10 times. The p value was calculated using two-sided Student’s t test. **d, e**, Scatter plots comparing the day 16 vs day 1 log2 fold change (LFC) values of cysteines in Common Essential dependencies in KMS26 versus PC14 cells. The identified essential cysteines (FDR cutoff: 10%, LFC cutoff: −0.6) are highlighted for (**d**) ABE and (**e**) CBE libraries in green and yellow, respectively. **f, g**, Dot plots comparing the dropouts of cysteines from Strongly Selective cancer dependencies using PC14 (**f**) or KMS26 (**g**) cells. Each pair of points compares the day 16 vs day 1 LFC for the same cysteine. The cysteines were included based on the first essentiality filter (FDR cutoff: 10%, LFC cutoff: −0.6) only using data from one cell line (**f**) PC14 or (**g**) KMS26 without considering the dependency difference between the two cell lines. A second selectivity filter (greater dropout in the more dependent cell line, LFC difference > 0.3) was then applied, and cysteines passing this filter are shown in red. Only Strongly Selective proteins with gene-level dependency scores that are sufficiently different (CERES> 0.8) between PC14 and KMS26 were included in the analysis. The p values were calculated using two-sided paired Student’s t test based on all cysteines that pass the first essentiality filter.

**Extended Data Fig. 3.**
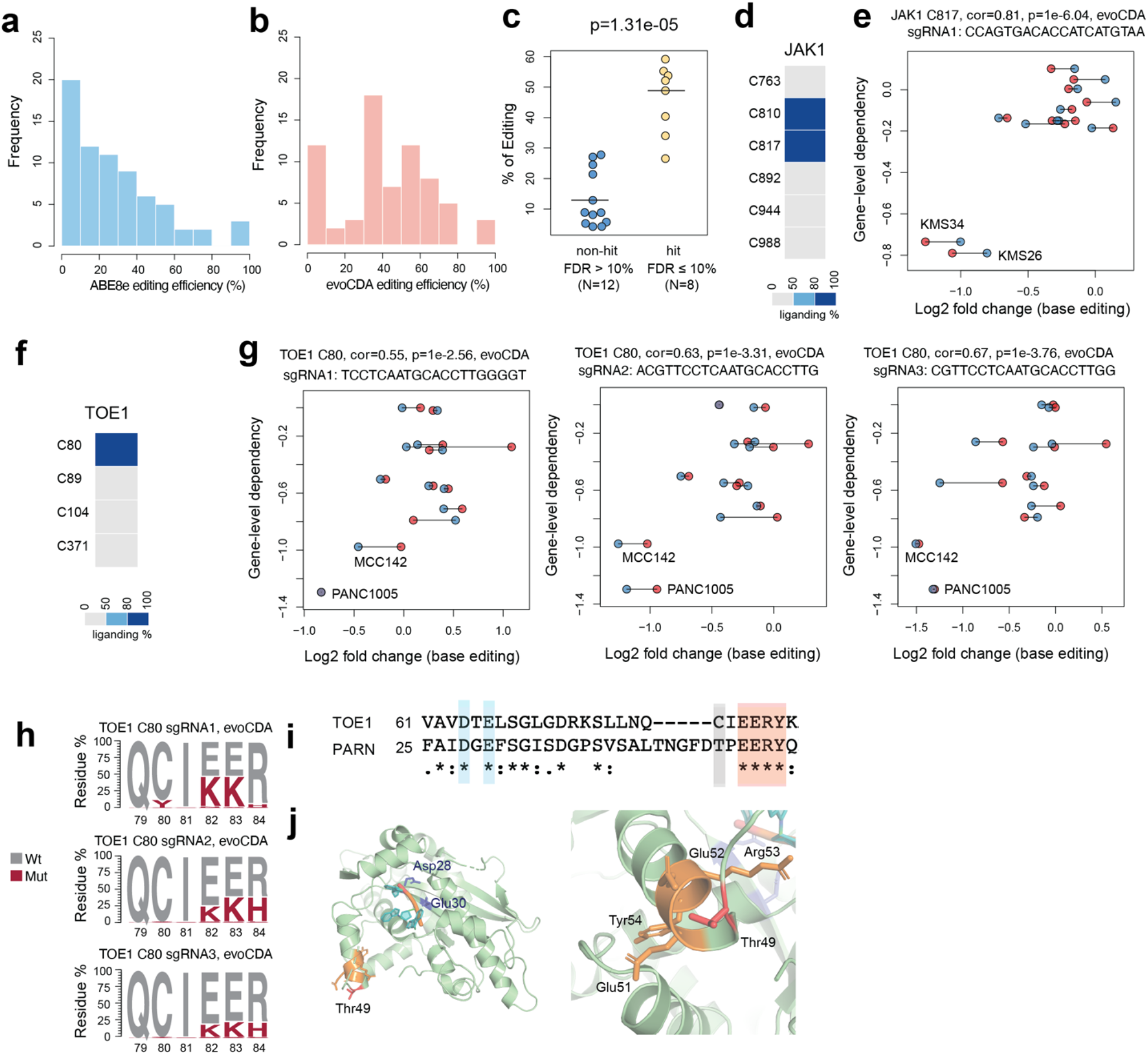
Base editing reveals essentiality of ligandable cysteine regions in Strongly Selective cancer dependency proteins. **a, b**, Histogram showing the gene editing efficiencies for 70 sgRNAs using (**a**) ABE8e-NG or (**b**) evoCDA-NG in PC14 cells. The relative residue mutation frequency five days after lentivirus infection was quantified using targeted genomic PCR and amplicon sequencing. The percentage of gene editing was estimated by the ratio of edited allele reads that correspond to missense mutations in the target region over total mapped reads in a quantification window of 20bp. Data represent average values of two independent experiments. **c**, Beeswarm plot comparing the editing efficiencies among the sgRNAs that did or did not create significant base editing-knockout correlations. Each point represents a different sgRNA for 20 total sgRNAs designed for FOXA1 or TOE1. The corresponding genomic sites were individually sequenced to quantify the editing efficiencies as described in **a, b** above. The p value was calculated using two-sided Student’s t test. **d**, The fragment electrophile ligandability profile for cysteines in JAK1, showing ligandability of C817. Note that C810 and C817 on the same tryptic peptide, so their ligandability cannot be distinguished by cysteine-directed MS-ABPP. However, other studies have verified that C817 is the liganded cysteine in JAK1^28^. **e,** Scatter plots for the JAK1_C817 region showing the correlation between base editing-induced cell line dropout data acquired herein and gene disruption-induced cell line dropout data in the Cancer Dependency Map. The two independent biological experiments for base-editing data in each of the twelve cell lines are shown in blue and red. **f**, The fragment electrophile ligandability profile for cysteines in TOE1, showing ligandability of C80. **g**, Scatter plots for the TOE1_C80 region showing the correlation between base editing-induced cell line dropout data acquired herein and gene disruption-induced cell line dropout data in the Cancer Dependency Map. The two independent biological experiments for base-editing data in each of the twelve cell lines are shown in blue and red. Each plot shows a different sgRNA. **h**, TOE1 amino acid mutagenesis generated by different sgRNAs and the indicated base editors. The relative residue frequency five days after lentivirus infection was quantified using targeted genomic PCR and amplicon sequencing. Data represent average values of two independent experiments. **i**, Alignment of TOE1 and PARN protein sequences. The conserved catalytic residues are highlighted in blue and the EERY sequence is highlighted in red. **j**, TOE1 homology model based on the crystal structure of PARN in complex with RNA (PDB: 2A1R). The TOE1_C80 corresponding residue PARN_T49 and the EERY sequence is highlighted.

**Extended Data Fig. 4.**
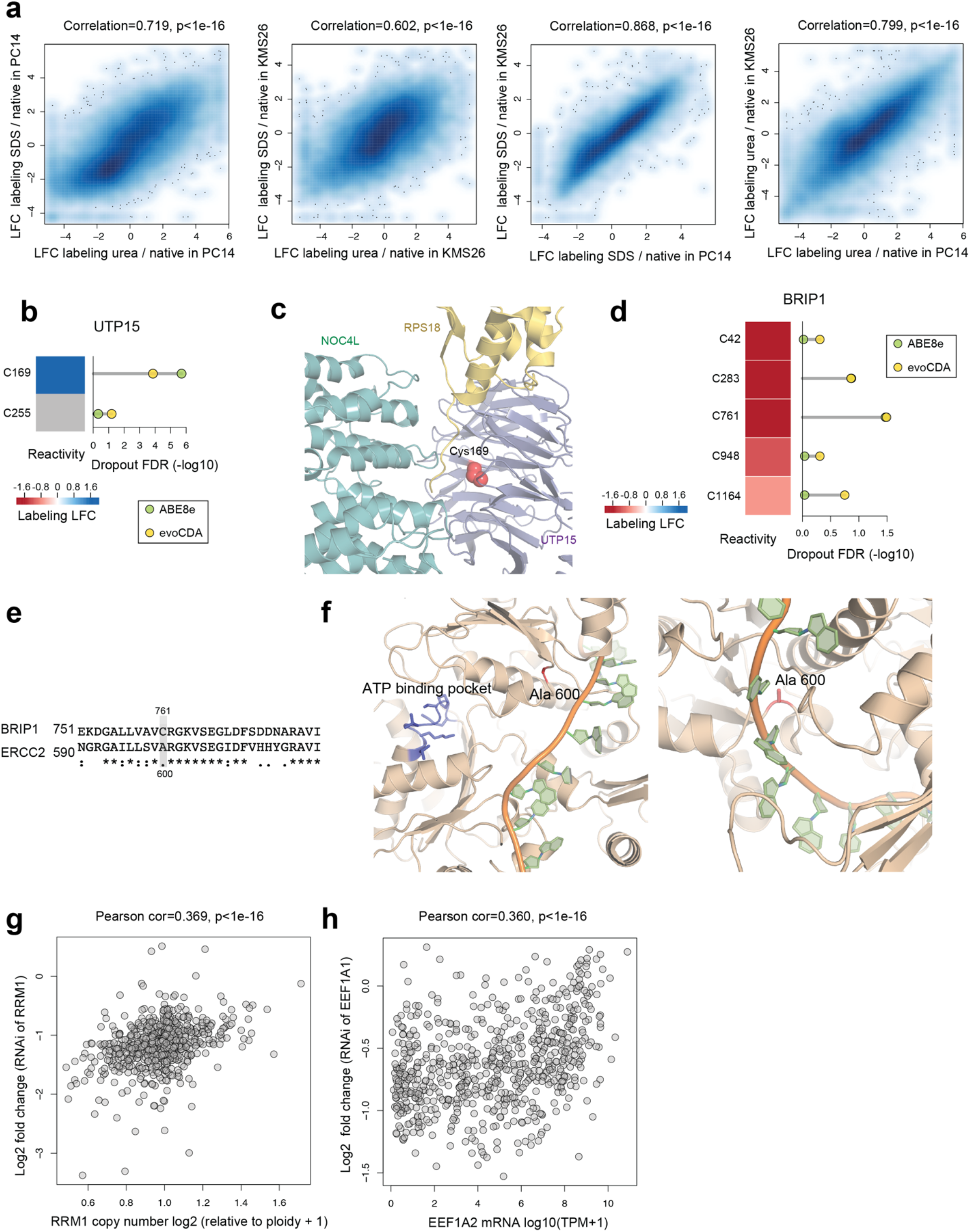
Prioritizing essential cysteines with ligandability potential by quantitative cysteine reactivity profiling in native and denatured proteomes. **a**, (from left to right) Density scatter plots comparing the effect of i) denaturants urea vs SDS on IA-DTB labeling in PC14 cells (left) and KMS26 cells (left middle); and ii) SDS (right middle) or urea (right) on IA-DTB labeling in PC14 vs KMS26 cells. Each point represents a quantified IA-DTB labeled cysteine. **b**, The reactivity in denatured/native proteomes (left) and significance of dropout (right) for cysteines in UTP15. **c**, Structure of UTP15 in complex with NOC4L and RPS18 as part of the human small subunit processome (PDB: 7MQ8). The side chain of essential, unreactive (buried) C169 is highlighted in red. **d**, The reactivity in denatured/native proteomes (left) and significance of dropout (right) for cysteines in BRIP1. **e**, Alignment of BRIP1 and ERCC2 protein sequences. **f**, BRIP1 homology model based on the crystal structure of ERCC2 in complex with DNA (PDB: 6RO4). The BRIP1_C761 corresponding residue ERCC2_A600 and the ATP-binding pocket is highlighted. **g**, Scatter plot showing the association between RRM1 gene copy number and sensitivity to RNA interference (RNAi)-dependent suppression of cancer cell growth. Each point represents a cancer cell line in the Cancer Dependency Map. **h**, Scatter plot showing the association between mRNA expression levels for the EEF1A1 paralog EEF1A2 and sensitivity to RNAi-mediated EEF1A1 knockdown-dependent suppression of cancer cell growth. Each point represents a cancer cell line in the Dependency Map. For **g** and **h**, p values were calculated using a two-tailed Pearson correlation test.

